# Acetazolamide reduces exercise capacity following a five-day ascent to 4559 m on Monte Rosa

**DOI:** 10.1101/105726

**Authors:** Arthur R. Bradwell, Kimberly Ashdown, Carla Gallagher, John Delamere, Owen D. Thomas, Samuel J.E. Lucas, Alex D. Wright, Stephen J. Harris, Stephen D. Myers, Birmingham Medical Research Expeditionary Society

## Abstract

Acetazolamide (Az) is widely used to prevent and treat the symptoms of acute mountain sickness (AMS) but whether it alters exercise capacity at high altitude is unclear. Az (250 mg twice daily) or placebo were administered to 20 healthy adults (age range, 21-77 years) in a double-blind, randomized manner. Participants ascended over five days to 4559 m, before undertaking an incremental exercise test to exhaustion on a bicycle ergometer, with breath-by-breath gas measurements recorded using a portable gas analysis system. Maximum power output (P_max_) was reduced on Az compared with placebo (*p*=0.03), as was maximum O_2_ uptake (VO_2max_) (20.7 vs 24.6 mL/kg/min; *p*=0.06) and maximum expired CO_2_ (VCO_2max_) (23.4 vs 29.5 mL/kg/min; *p*=0.01). Comparing individuals matched for similar characteristics, Az-treated participants had smaller changes than placebo-treated participants in minute ventilation (88 vs 116 L/min: *p*=0.05), end tidal O_2_ (6.6 vs 9.3 mm Hg: *p*=0.009), end-tidal CO_2_ (−2.3 vs −4.2 mm Hg: *p*=0.005), VO_2max_ (9.8 vs 13.8 mL/kg/min; *p*=0.04) and VCO_2max_ (14.7 vs 20.8 mL/kg/min; *p*=0.009). There was a negative correlation between the mean ages of paired vs placebo-treated individuals and differences in P_max_ reductions from base-line to altitude (*r* =−0.83: *p*<0.005) and HR_max_ at altitude (*r*=−0.71; *p*=0.01). Glomerular filtration rate (measured at sea-level) declined with increasing age (*r*=−0.69; *p*=0.001). Thus, 250mg of Az twice daily reduced exercise performance, particularly in older individuals. The age-related effects of Az may reflect higher tissue concentrations due to reduced drug clearance in older people.

## New and noteworthy

We have identified a reduction in exercise performance (maximum power output and VO_2max_) at high altitude in individuals given commonly prescribed doses of acetazolamide for acute mountain sickness. This reduction was greater in older people (>50 years) possibly due to reduced renal clearance of the drug. Results indicate that it may be appropriate for older people to use smaller doses of acetazolamide at altitude.

## Introduction

Acetazolamide (Az) is an important medication for the prevention of acute mountain sickness (AMS). This was demonstrated initially by Forwand et al, (1968) followed by a large study on Everest trekkers (Hackett et al. 1976) with an accompanying editorial (‘See Nuptse and Die’ - Rennie 1976). In controlled studies, Az has been shown to increase arterial oxygen saturations at all altitudes and to assist in ascents to Everest Base Camp, the summit of Kilimanjaro, and elsewhere (Bradwell et al, 1981; Greene et al, 1981; Stokes et al, 2010). A recent meta-analysis of 24 placebo controlled trials comparing 1011 Az-treated with 854 placebo treated individuals showed convincing evidence of its value for the prevention of AMS (Keyser et al, 2012). Escalating doses of Az from 250 mg, to 500 mg and 750 mg per day reduced AMS symptoms by 45%, 50% and 55% respectively.

An important but controversial aspect of Az use is its effect on exercise at altitude. Of five chamber studies examining the impact of Az during short-term exposure to hypoxia, three showed reduced VO_2max_ and/or time to exhaustion (McLellan et al, 1988; Stager et al, 1990; Garske et al, 2003), one slight increase in VO_2max_ (Schoene et al, 1983) and one no effect (Jonk et al, 2007). Similar inconsistent findings have been reported in natural high-altitude environments. Hackett et al. (1985) gave Az acutely to well-acclimatised individuals and observed a reduction in exercise performance (time at maximum workload), whereas Faoro et al, (2007) observed no effect. When Az is used prophylactically, a common practice among trekkers, results are again inconsistent. After a 14-day trek to 4846 m, VO_2max_ and exercise endurance were greater on Az compared with placebo (Bradwell et al, 1986). Conversely, when used during early acclimatisation (18-24 hours), Az reduced exercise performance, but only in individuals over 50 years of age (Bradwell et al, 2014).

Surveys of trekkers add no further enlightenment; Az is widely used but there are no reported adverse effects on exercise. Presumably, those affected would ascend more slowly and ascribe any perceived weakness to altitude or poor fitness. Nevertheless, exercise capability is an important issue when there are time-restricted climbing schedules. For instance, on Kilimanjaro 35% of climbers use Az yet the overall summit success rate can be as low as 50-60% on three and four day ascents (Stokes et al, 2010).

There are also questions about the use of Az in older individuals. Because Az is excreted un-metabolised in urine (Heller et al, 1985; Chapron et al, 2000), drug levels increase with reduced glomerular filtration rates. Given the known age-related decline in renal function, Az concentrations might increase sufficiently to induce excess acidosis, thereby inhibiting exercise. Because increasing numbers of older people are exploring high mountains (Saito et al, 2002), questions regarding the effect of Az on exercise need to be answered, particularly as older people may be less susceptible to AMS (Honigman et al, 1993). The purpose of this study was to assess the effects of Az on exercise performance in young and older individuals during a typical alpine high-altitude climb.

## Materials and methods

*Subjects.* Twenty healthy individuals were recruited (16 male, four female). Ages ranged from 21 to 77 years; 13 were under 26 years, with six males over 50 years. Fifteen individuals had previous experience of high altitude and their susceptibility to AMS was categorised into mild or moderate. No participant had suffered from high-altitude pulmonary or cerebral edema (severe AMS), resided above 1,500 m in the previous 2 months, taken any relevant medication, or undertaken any intense physical activity for 7 days before baseline testing. All participants had free access to fluids with no measurement of hydration status and all were non-smokers.

Individuals were paired for similar characteristics according to the following hierarchy: (1) age, (2) sex, (3) previous AMS susceptibility, and (4) body mass. Since all participants were fit and healthy, they were well matched for exercise performance using these four criteria (Table 1). Each of the pair took either Az 250 mg twice daily or placebo (lactose powder) in capsule form, following a double-blind design. Medication was commenced the day before altitude exposure for a total of ten doses over 5 days. Because of potential un-blinding linked to Az side-effects, individuals were requested not to discuss any aspects of their medication with others, including the investigators.

**Table 1.** Participant characteristics (mean ± 1SD) at base-line (150 m).

|  | Placebo (10) | Acetazolamide (10) |
| --- | --- | --- |
| Gender: male/female | 8/2 | 8/2 |
| Age (range of years) | 33 ± 17 (21-66) | 40 ± 23 (21-77) |
| Height (cm) | 176 ± 8 | 177 ± 11 |
| Weight (kg) | 73 ± 13.6 | 73 ± 13 |
| AMS history: mild:moderate:unknown | 3: 4: 3 | 6: 2: 2 |
| VO<sub>2max</sub> (mL/kg/min) | 40.3 ± 6.3 | 35.9 ± 6.1 |
| Maximum power output (W) | 234 ± 51 | 215 ± 39 |
| Heart rate maximum (beats/min) | 185 ± 20 | 169 ± 26 |

There were no significant differences (all $p > 0.05$ ) between groups (Mann-Whitney U test).

*Baseline experiments and equipment.* Two weeks before ascent, graded baseline exercise tests were carried out in Birmingham (150 m above sea-level) to determine maximum power output (P_max_) and maximum oxygen uptake (VO_2max_). Exercise tests were performed on a light-weight (25kg), recumbent bicycle ergometer designed for altitude studies (Alticycle) as previously described (Bradwell et al, 2014). This was used because it is readily portable and since head and spine are stationary, breathing masks and cardiopulmonary monitoring sensors are held reliably in place.

After a 5-minute, self-paced warm-up, participants commenced at 50-100 W depending on their self-reported level of fitness. Using a cadence of 60 rev/min, power was increased every three minutes by 25 W or 15 W, for men or women respectively, until 80% of the predicted maximum. Thereafter, power steps were increased every minute until volitional exhaustion. Breath-by-breath gas measurements were recorded of minute ventilation (VE), end-tidal oxygen (PetO_2_) and carbon dioxide (PetCO_2_) concentrations, oxygen uptake (VO_2_), and expired CO_2_ (VCO_2_) using a Cosmed K4b2 (Cosmed, Rome, Italy) (Bassett et al, 2001). Heart rate (HR), peripheral blood O_2_ saturation (SpO_2_) and perceived exertion were recorded at rest and for every exercise stage. The maximum VO_2_ attained (VO_2max_) was determined as the highest 20-second moving average in VO_2_, together with the corresponding P_max_.

To assess the exercise intensities appropriate at altitude, two individuals were tested at 50%, 60% and 70% of their predicted VO_2max_ in a normobaric hypoxic chamber (TISS Model 201003-1, TIS Services, Mestead, UK) at an inspired oxygen fraction (FI_O2_) equivalent to 4559 m (the altitude of the Margherita Hut on Monte Rosa). These individuals were able to sustain 60% of their sea-level maximum for three minutes, which is consistent with previous reports of power reduction at altitude (Fulco et al, 1998). This was used as the likely aerobic maximum for each participant at altitude. In addition, blood samples were taken in Birmingham for creatinine measurements in order to calculate glomerular filtration rates using the Cockcroft-Gault equation (Froissart et al, 2005).

*Ascent profile and altitude studies.*
Figure 1 shows the ascent profile for Monte Rosa used in this study compared with typical acclimatization and mountaineering gradients for Mt Blanc and Kilimanjaro (~35,000 summit attempts each per year).

**Figure 1.**
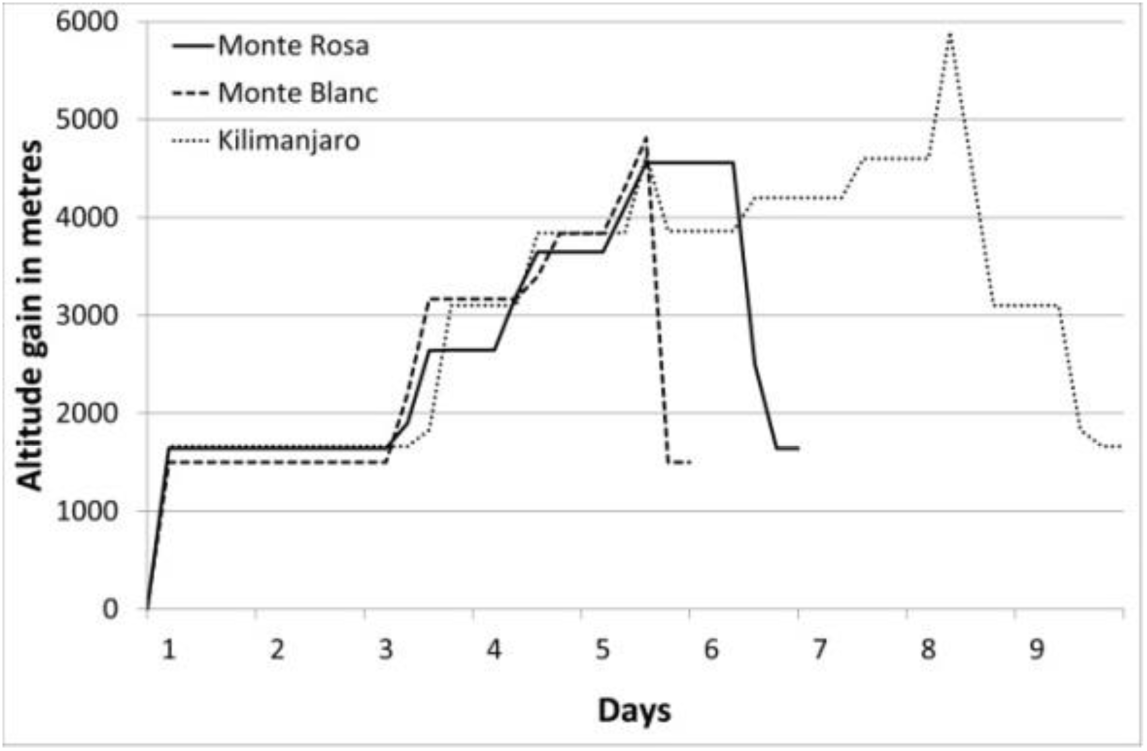
Comparison of the daily altitudes achieved on Monte Rosa (4559 m) compared with typical ascent profiles for Mt Blanc (4810 m) and Kilimanjaro (5895 m)

Individuals travelled overland and by sea from the UK to Gressonay (1640 m) in Italy over an 18-hour period, followed by a two-night stay that included an acclimatization ascent to 2800 m. Free access to fluid was allowed and hydration status was not measured. All participants ascended on sequential days to 2646 m, 3647 m and 4559 m. Matched pairs ascended together and at their own pace. Exercise tests commenced at the Margherita Hut shortly after arrival. Self-assessed questionnaires using the Lake Louise (LL) scoring system for AMS (Roach et al, 1993) were recorded every morning and evening. Since the exercise tests were carried out at different times of the day, the mean of the morning and evening scores were considered to be the best assessment of AMS. Individuals were asked immediately before their exercise test which medication they thought they were taking.

At altitude, exercise tests were performed in a similar manner to baseline. Following resting measurements, participants completed a 5-minute self-paced warm-up. The test commenced at the power output equivalent to 23% of each participant’s baseline P_max_ and was increased by 7-8% every three minutes up to 60% of baseline P_max_ and thereafter every minute until volitional exhaustion.

*Statistics.* Statistical analyses were performed using SPSS version 21 (IBM Corp, Portsmouth, UK). Mann-Whitney U test was used to analyse differences of means; Pearson’s chi-squared test was used to analyse observed differences between matched pairs and Pearson’s correlation.

*Ethics.* The study was approved by Chichester University Research Ethics Committee (protocol number: 1314_42) and was performed according to the Declaration of Helsinki. All individuals gave signed informed consent.

## Results

*Baseline studies.* Results of the baseline tests are shown in Table 1. There were no significant differences between placebo- and Az-treated groups.

*Altitude studies.* Daily ascents were completed as planned and comprised 3-5 hours of moderate to strenuous exercise. At the Margherita Hut (4559 m), exercise tests were commenced 2-3 hours after arrival and completed by 10 hours. Matched pairs of placebo- and Az-treated individuals were assessed within 90 minutes of each other. Questionnaires recorded immediately before the exercise test (but analysed later) indicated that 11 had correctly guessed their medication, seven did not know, while two on placebo thought they were taking Az. Mean AMS symptom scores for each group are shown in Figure 2. Initial marginally elevated scores were after the 18-hour overnight bus journey and showed increased levels of sleeplessness. On the day of the exercise test, summated morning and evening Lake Louise AMS scores were similar in the two groups (Az vs placebo: 4.2 [±4.1] vs 4.2 [±2.7]; *p*=0.7).

**Figure 2.**
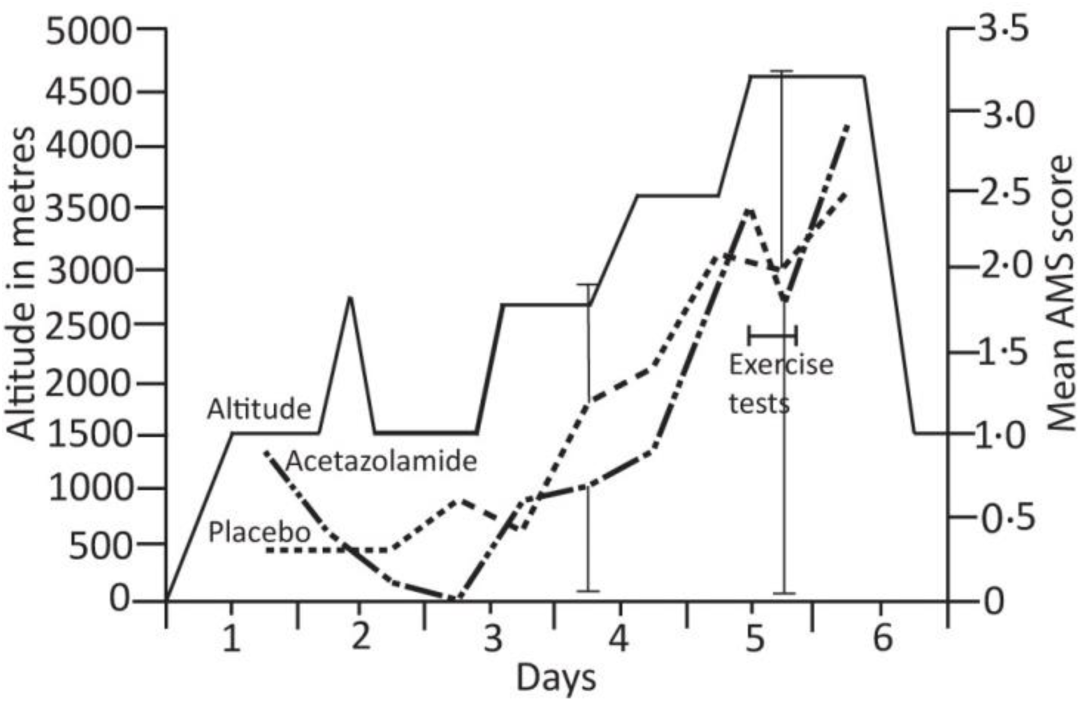
Twice daily Lake Louise AMS scores of the placebo and Az groups alongside the daily ascent profile. No significant differences were seen between the two groups at any time point. Two representative standard deviation bars are shown.

The exercise test results are shown in Table 2 and Figures 3–6. At rest, VO_2_, VCO_2_ and respiratory exchange ratios were similar in both groups while resting SpO_2_ and PetO_2_ were higher in the Az-treated group and PetCO_2_ was lower.

**Table 2.** Altitude performance (4559 m) at rest, 45%, and maximum power (P_max_) for placebo and Az groups, and comparisons between matched pairs.

| Test variable | Placebo vs Az at rest <sup>^</sup> | Placebo vs Az at 45% $P_{\max}$ <sup>^</sup> | Placebo vs Az at $P_{\max}$ <sup>^</sup> | Comparison of placebo vs Az for matched pairs | |
| --- | --- | --- | --- | --- | --- |
| | | | | Change from rest to $P_{\max}$ <sup>*</sup> | Difference in pairs at $P_{\max}$ <sup>+</sup> |
| $P_{\max}$ (W) as % of sea-level values | - | - | #76.6 (7.4) v 65 (14.1)<br>**$p=0.03$** | - | **$p=0.007$** |
| Heart rate (b/min) | 92(18.4) v 93(15.7)<br>$p=0.88$ | 146 (17) v 140 (17)<br>$p=0.43$ | 167 (16) v 154 (25)<br>$p=0.33$ | 75 (21) v 61 (20)<br>$p=0.1$ | **$p=0.007$** |
| RPE (Borg scale: 6-20) | 10(2) v 9(1)<br>$p=0.36$ | 14(2) v 14(1)<br>$p=0.56$ | 17(2) v 17 (2)<br>$p=0.81$ | 7 (2) v 8 (2)<br>$p=0.36$ | NS |
| VE (L/min) | 34(9) v 37(10)<br>$p=0.48$ | 75(13) v 88(22)<br>$p=0.11$ | 150(22) v 125(27)<br>**$p=0.04$** | 116 (21) v 88 (28)<br>**$p=0.05$** | NS |
| SpO<sub>2</sub> (%) | 79.0(4.1) v 84.6(4.7)<br>**$p=0.004$** | 74.1(5.2) v 75.5(5.1)<br>$p=0.55$ | 77.0(4.4) v 77.2(6.3)<br>$p=0.7$ | -2.4 (5.7) v -7.4 (6.3)<br>$p=0.13$ | NS |
| PetO<sub>2</sub> (mm Hg) | 54.9(2.7) v 58.4(2.7)<br>**$p=0.007$** | 57.8(2.7) v 61.9(3.8)<br>**$p=0.01$** | 64.2(2.0) v 65.0(2.0)<br>$p=0.45$ | 9.3 (1.8) v 6.6 (2.3)<br>**$p=0.009$** | $p=0.07$ |
| PetCO<sub>2</sub> (mm Hg) | 21.0(1.5) v 17.5(1.5)<br>**$p=0.0006$** | 20.5(2.1) v 16.6(2.7)<br>**$p<0.01$** | 16.8(1.5) v 15.2(1.5)<br>**$p=0.05$** | -4.2 (1.37) v -2.3 (0.95)<br>**$p=0.005$** | **$p=0.0003$** |
| VO<sub>2</sub> uptake: (mL/kg/min) | 10.8 (3.2) v 10.9 (2.8)<br>$p=0.97$ | 19.5 (5.5) v 18.4 (2.8)<br>$p=0.56$ | 24.6 (5.1) v 20.7 (5.2)<br>$p=0.063$ | 13.8 (4.9) v 9.8 (6.2)<br>**$p=0.04$** | **$p=0.0003$** |
| VCO<sub>2</sub> production (mL/kg/min) | 8.7(2.5) v 8.7(2.3)<br>$p=0.91$ | 18.7(4.8) v 18.3(2.3)<br>$p=0.82$ | 29.5(4.6) v 23.4(5.5)<br>**$p=0.01$** | 20.8 (5.0) v 14.7 (6.0)<br>**$p=0.009$** | **$p=0.0003$** |
| VE/VO<sub>2</sub> produced | 44(5.8) v 50(6.8)<br>**$p=0.04$** | 55.0(8.0) v 70.8(18.2)<br>**$p=0.03$** | 86(16.0) v 88(11.5)<br>$p=0.78$ | 42 (13.0) v 38 (7.1)<br>$p=0.68$ | NS |
| VE/VCO<sub>2</sub> produced | 55(4.1) v 64(5.3)<br>**$p=0.001$** | 56.9(6.7) v 70.8(14.2)<br>**$p=0.02$** | 70(6.3) v 79(9.7)<br>**$p=0.04$** | 15 (5.2) v 15 (4.8)<br>$p=0.80$ | **$p=0.0003$** |
| CO<sub>2</sub> prod<sup>n</sup>/O<sub>2</sub> uptake (RER) | 0.81(0.07) v 0.78(0.06)<br>$p=0.44$ | 0.96(0.06) v 0.99(0.07)<br>$p=0.44$ | 1.23(0.16) v 1.12(0.1)<br>$p=0.14$ | 0.42 (0.12) v 0.34 (0.09)<br>$p=0.19$ | **$p=0.007$** |

Data are mean values ( $\pm 1$ SD) with $p$ -values in bold when $p=0.05$ or less. Symbols indicate type of statistical test used: <sup>^</sup> Mann-Whitney U test for differences of means; <sup>\*</sup> Pearson's correlation for changes in paired individuals; <sup>+</sup> Pearson's chi-squared test for observed differences between matched pairs; <sup>#</sup> % reduction of individuals mean values between base-line and altitude. RPE – rate of perceived effort; VE – ventilation; SpO<sub>2</sub> – peripheral blood O<sub>2</sub> saturation; PetO<sub>2</sub> – end tidal O<sub>2</sub>; PetCO<sub>2</sub> – end tidal CO<sub>2</sub>; VO<sub>2</sub> – O<sub>2</sub> uptake; VCO<sub>2</sub> – CO<sub>2</sub> production; RER – respiratory exchange ratio.

During exercise, all participants were able to achieve 45% of their baseline performance. At this comparable time point, VE and PetO_2_ were on average higher, while PetCO_2_ was lower in the Az group compared with the placebo group (Table 2 and Figure 4). Thereafter, greater power was generated by placebo participants both as a group (*p* = 0.03) (Figure 3) and in the matched pairs (*p* = 0.007) (Table 2).

**Figure 3.**
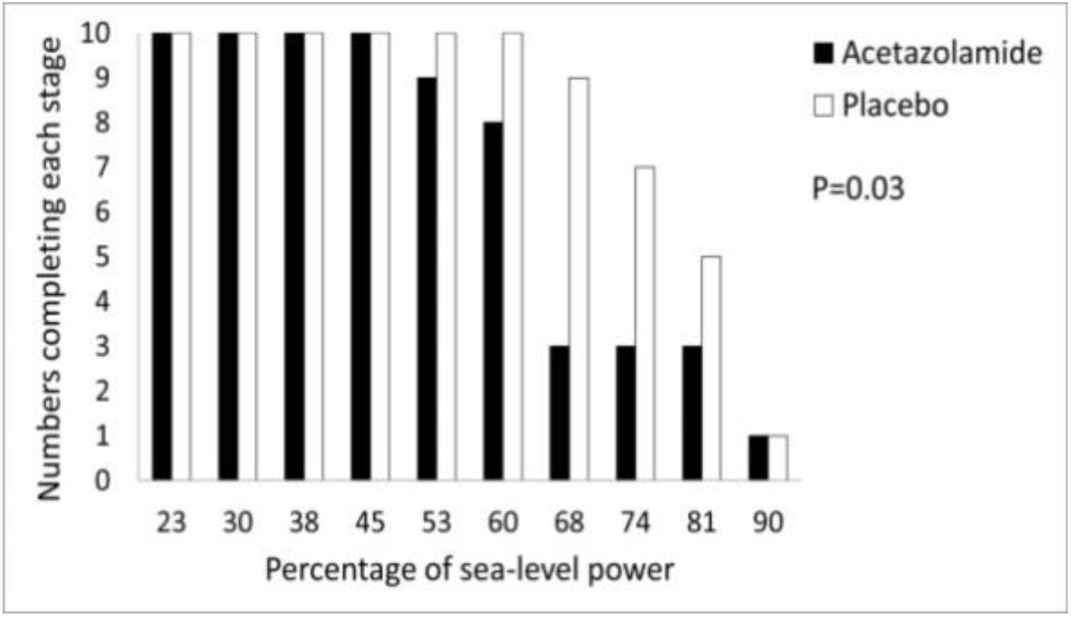
Percentage of sea-level power achieved at 4559 m by participants in each treatment group.

As exercise intensity increased, PetO2 increased while SpO_2_ and PetCO_2_ reduced (Figure 4).

**Figure 4.**
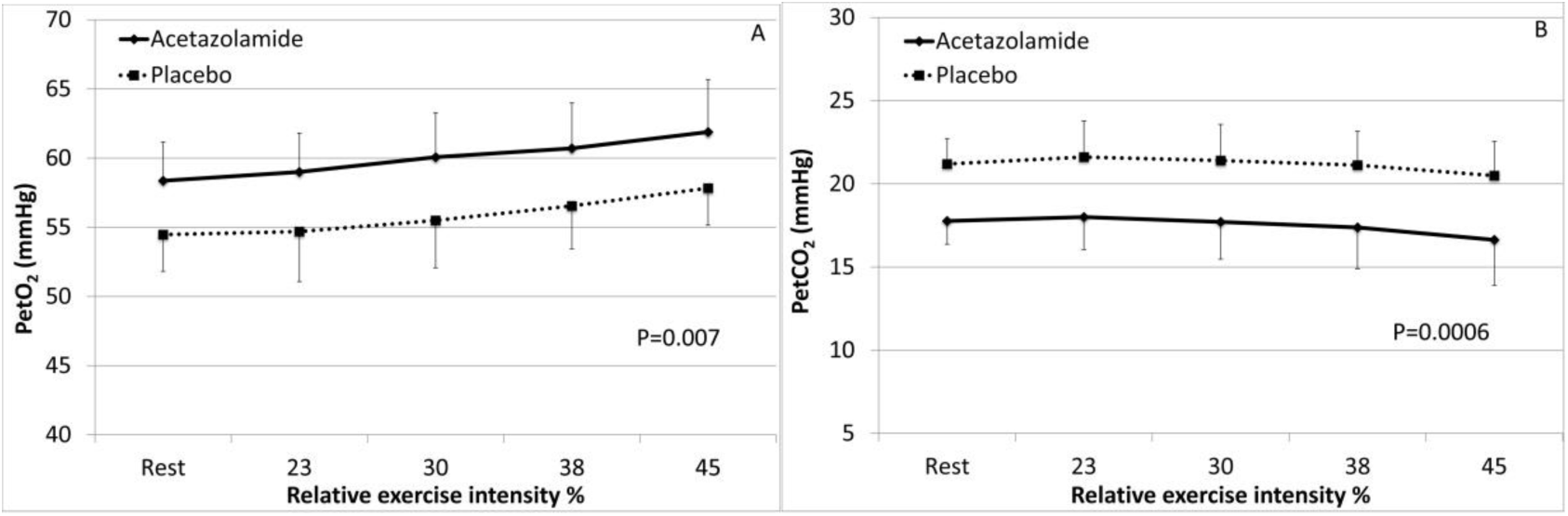
(A) PetO2 and (B) PetCO2 in placebo and Az groups with increasing exercise intensities to 45% of baseline values (plus SDs). Exercise intensities beyond 45% are not shown due to the progressive failure of individuals to complete each stage, which created unbalanced groups at higher intensities (Figure 3).

Comparison of individual results show that those on placebo had similar heart rates and VO_2_ at rest to Az individuals, but higher values at P_max_ (Figure 5A and C) although differences did not reach statistical significance for heart rate data. However, comparison of matched pairs at P_max_ demonstrated higher values in the placebo-treated individuals (Figures 5B and D).

**Figure 5.**
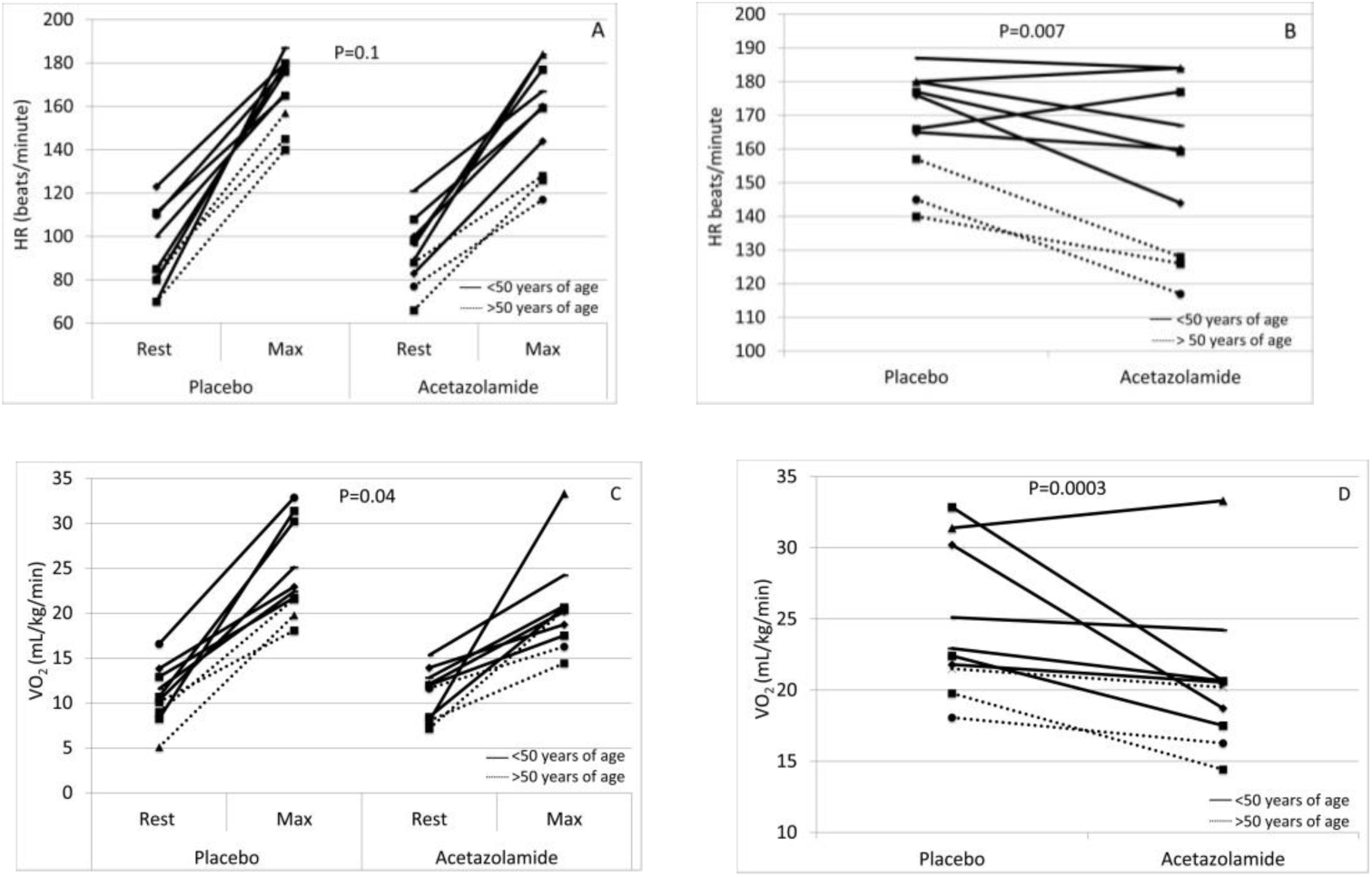
Altitude studies at 4559 m showing heart rate (HR) and oxygen uptake (VO_2_) at rest and P_max_ for each participant (A and C) and comparison of matched pairs for HR and VO_2_ at P_max_ (B and D). Individuals and matched pairs >50 years of age are shown as dotted lines.

Correlation analysis of the matched pairs was used to assess the relationship between age and the effect of Az on the physiological parameters measured. There was a negative correlation between the mean ages of placebo- and Az-treated pairs and differences in HRmax at altitude (*r* = −0.71, *p* = 0.01) (Figure 6A). Similarly, there was a negative correlation between the mean ages of pairs and the size of the reduction in P_max_ (base-line to altitude) between placebo- and Az-treated individuals (*r* = −0.83: *p*<0.005) (Figure 6B). No other differences between pairs significantly correlated with the mean ages of the paired individuals.

**Figure 6.**
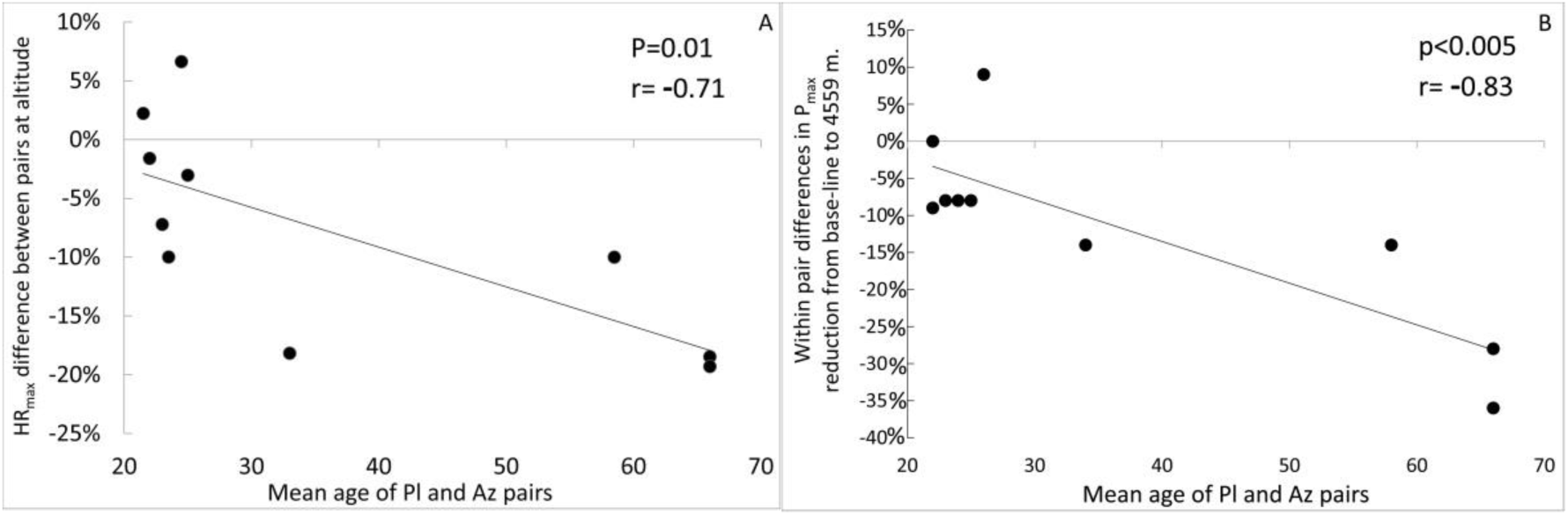
P_max_ at altitude showing the relationship within paired participants (placebo vs Az). Comparison of mean ages and (A) differences in the maximum heart rate (HR_max_), and (B) differences in reduction in maximum power (P_max_) from sea-level (Pearson’s correlation).

There was a negative correlation between eGFR of participants and age (*r*=−0.69; *p*=0.001) (Figure 7).

**Figure 7.**
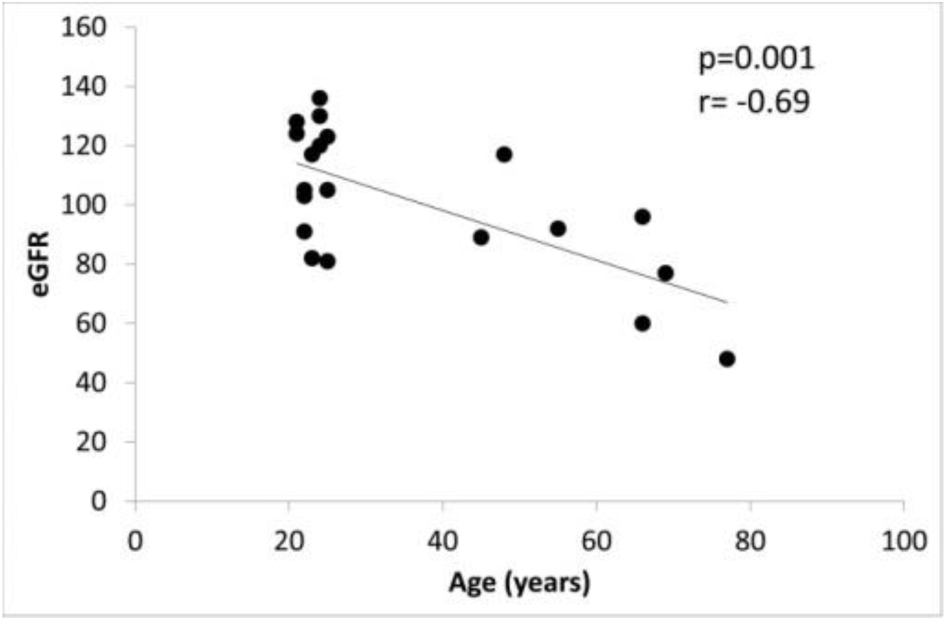
Relationship between ages of participants and estimated glomerular filtration rate (eGFR).

## Discussion

The main finding of this study was the greater reduction in P_max_ at altitude in individuals taking Az compared with those on placebo, with the effect more pronounced in those over 50 years (Figure 6B). We observed that, although Az stimulated respiratory centres leading to higher SpO_2_ and PetO2 at rest, this did not translate into an improvement in exercise performance. Indeed, during the incremental increases in exercise intensity, SpO_2_ fell more in the Az group while VO_2_ (Table 2 and Figure 5) and VCO2 (Table 2) showed larger increases in the placebo group, indicating a greater exercise capacity. At altitude, heart rates were similar in both groups at rest, but Az-treated participants showed smaller increases at P_max_. This was apparent in the matched pairs (Figure 5B) and particularly in older individuals (Figure 6A).

Our finding of impaired P_max_ on Az is consistent with a previous mountain study at lower altitude (3459 m) (Bradwell et al, 2014) and a detailed chamber study at a simulated altitude of 4200 m (Garske et al, 2003). The latter study also reported reduced blood pH at near-peak exercise on Az. Such pH changes reflect the slowed CO2 excretion kinetics and the renal-metabolic acidosis mediated by Az, via inhibition of carbonic anhydrase (Scheuermann et al, 1999). Acidosis within muscle cells leads to decreased endurance time during exercise and inhibition of glycolytic enzymes, especially phosphofructokinase (Jones et al, 1997). Thus, impaired CO_2_ elimination is likely to have contributed to acidosis in our Az group leading to reduced P_max_.

The nine studies that have examined Az effects on exercise performance do not all agree with these observations. This may be explained, in part, by differences in altitude/FIO2 attained, Az dose, ages of the participants, nature of control groups and exercise performance measures that were assessed. Furthermore, some studies were chamber simulations of altitude while others were carried out in natural high-altitude environments. Of the five chamber studies, one showed a slight increase in exercise performance (Schoene et al, 1983) another found no effect (Jonk et al, 2007) while three others showed reductions in measures of exercise performance (Garske et al, 2003; McLellan et al, 1988; Stager et al, 1990). The latter two studies used exercise endurance as the main outcome measure, which can be strongly influenced by motivation (Fulco et al, 1998), while Garske et al. (2003) used the more objective VO_2_max test.

The four field-based studies at altitude have also produced conflicting results. Hackett et al. (1985) and Faoro et al. (2007) both used 750mg of Az in fully acclimatized individuals over a 24-hour period before the exercise test. However, the studies were conducted at different altitudes (6300 m and 4700 m) and produced different results, one showing reduced VO_2_max (Hackett et al, 1985) and the other no effect (Faoro et al, 2007). Older people (>40 years) were included in both studies. However, Hackett et al. studied only two older individuals on Az, and while Faoro et al. studied 15 older individuals there was no indication of whether age or male/female ratios were well balanced in the treatment and control arms. Our group conducted the two other field studies reported to date (Bradwell et al. 1986 and Bradwell et al. 2014), in which participants took Az prophylactically at a dose of 500mg per day. In our earlier study (Bradwell et al. 1986), which contained no older individuals, participants were studied after 10 days trekking followed by three to four days at 4846 m before exercise testing. We observed an improved exercise endurance and higher VO_2_max in those on Az. We reasoned that this was due to reduced AMS in individuals on Az which meant that they had exercised and eaten more during two weeks at altitude leading to measureable retention of muscle mass and better performance during the exercise test. In contrast, our more recent study (Bradwell et al. 2014) was in partially acclimatized people who were tested after ascending rapidly to 3459 m. This was the first report to contain a high proportion of individuals over 50 years (nine out of 20) and showed that older individuals were less likely to complete a submaximal power test when taking Az. This is consistent with the results of our current study when we used a progressive exercise test to P_max_, combined with detailed metabolic data.

One important difference between studies is the dose of Az, which might explain some of the test result disparities. Larger doses are likely to have the greatest inhibiting effect on exercise ability, as shown in the data of Garske et al. (2003) who used 1000 mg per day for 2.5 days prior to testing. Since Az has a half-life of 8-12 hours (Ritschel et al, 1998) it would accumulate to produce a greater effect in those taking the drug for several days rather than only 24 hours (as is typically used in chamber studies). Az causes a metabolic acidosis only when sufficient bicarbonate has been excreted, which typically takes 24 hours or more (Burtscher et al, 2014). At altitude, this provides a beneficial ventilatory stimulus that opposes and limits the braking effect of hypocapnia on the full ventilatory response to hypoxia. In addition, carbonic anhydrase is also inhibited in red blood cells, brain, pulmonary and systemic vasculature, and muscles including the heart. The latter organ has high concentrations of carbonic anhydrase IV, which is 3.5 times more sensitive to Az inhibition than CAII and is 3 times more active in HCO_3_**^-^** dehydration (Baird et al. 1997). Furthermore, CAIV is present at high concentrations in the heart alongside CAIX and CAXIV (Waheed and Sly 2014). This suggests it has important cardiac functionality. As illustrated in Figure 6A, heart rate failed to increase as much in the Az group during exercise, particularly in the older participants. Age was also associated with reduced P_max_ in individuals on Az (Figure 6B). Since a reduced maximum heart rate would strongly influence P_max_, Az inhibition of cardiac carbonic anhydrase IV may partially account for the observed exercise limitation. Skeletal muscle, on the other hand, contains predominantly carbonic anhydrase III which is minimally inhibited by Az (Sly and Hu 1995).

The greater effect of Az on exercise performance in older people, raising interesting questions regarding the underlying mechanisms and dosage implications for this age group, especially given the increasing numbers of older people trekking at altitude (Saito et al, 2002). Aging leads to reduced VO_2_max, skeletal muscle mass (Conley et al, 2000) and renal function (Wetzels et al, 2007), all of which are directly affected by Az (Leaf and Goldfarb 2007; Swenson and Teppema 2007).

Since Az is largely excreted unchanged in urine (90% of an oral dose is excreted within 24 hours), tissue concentrations are dependent upon renal function (Chapron et al, 1989). However, there is an age-related decline in glomerular filtration rate, which falls by up to 10% per decade above 40 years of age (Wetzels et al, 2007). This causes Az to accumulate more in the tissues of older people such that toxicity is frequently observed when it is used for treating glaucoma, a disease of older people. Hence, patients with reduced glomerular filtration rates or on dialysis are prescribed doses of 125 mg per day or less (Chapron et al, 1989). In such patients, fatigue and lethargy are well-known side-effects. Consistent with this, we observed a strong negative correlation (*r* = −0.69; *p*=0.001) between glomerular filtration rate and age, supporting the notion that the older individuals taking Az in the current study were retaining more Az leading to higher tissue concentrations. This might have reduced cardiac function in addition to effects on other exercising muscles.

Finally, since Az is a mild diuretic, dehydration might have contributed to reductions in exercise performance in this group (Castellani JW et al. 2010). Hydration status was not measured but would be worth assessing in future high-altitude exercise studies.

*Implications.* Conventionally, Az usage at altitude is perceived as having few consequential side-effects (limited to minor taste disturbance and paresthesiae) or dosage implications. However, while higher dosing schedules provide additional benefits by reducing AMS symptoms (Keyser et al, 2012), this may be offset by reduced exercise performance. The narrow therapeutic window needs to be considered when Az is used for the prophylaxis of AMS, particularly in older people. Studies using doses as low as 125mg per day in older people are required with assessments of AMS scores, SpO_2_ and exercise performance. Furthermore, the results indicate that Az would not enhance exercise performance if taken by healthy individuals at altitude, but might have the opposite effect. As pointed out recently, age, body mass and gender need to be considered when prescribing Az to prevent AMS (Keyser et al, 2012).

*Limitations*. The investigation contained only 20 individuals so was relatively low-powered statistically, particularly for older participants. Further work is required to establish more clearly the age-related effect of Az on exercise performance. Furthermore, this investigation was a matched-pair rather than cross-over study, since the latter is difficult to achieve in a mountain environment because of acclimatization. We chose 500mg per day of Az in order to be comparable with many published studies; however, a commonly recommended dose for trekking is 250mg per day (Keyser et al, 2012), which may have less impact on exercise performance. Another limitation was the differing periods of altitude exposure during the study day. This was because people arrived in groups but testing was performed sequentially. To minimise any effect, paired individuals were tested within 90 minutes of each other. Measures of Az concentrations in the blood, blood and urine pH and ventilatory responses were not obtained and are likely to be relevant. We acknowledge that a VO_2_max test is only one measure of performance and other determinants of exercise performance and capacity at high altitude may be relevant in this context (e.g. endurance). Finally, exercise performance using a horizontal bicycle ergometer is less than that on a vertical bicycle; however, all participants were studied in the same manner at baseline and at high altitude. Since results were based upon changes in performance from rest to VO_2_max, we believe they would be similar using a vertical bike.

## Conclusion

We have shown that during an Alpine climb over five days, 500mg of Az daily reduced the ability to exercise to maximum, particularly in older people, while having no effect on reducing features of AMS. Although Az increased SpO_2_ and PetO_2_ at rest, O_2_ uptake and CO_2_ production were lower at P_max_ than those on placebo. The greater impact on exercise performance in older people probably reflected relative over-dosage of Az due to reduced renal clearance. While Az has value in preventing AMS, age-appropriate dosages (e.g. 250mg or even 125mg per day for people aged over 60) may be necessary to compensate for age-related changes in renal function and Az clearance. Clinical and laboratory studies are needed to evaluate lower-dose regimens in older individuals.

## Acknowledgments

The study was supported by the Birmingham Medical Research Expeditionary Society and the JABBS Foundation, that are both registered UK charities. Statistical analysis was undertaken by Dr Stephen Harding, Binding Site Ltd, Birmingham, UK. We thank the participants for their efforts in the exercise test.

### Conflict of interest statement

The authors declare that the research was conducted in the absence of any commercial or financial relationships that could be construed as potential conflicts of interest.

